# Crowding and binding: Not all feature-dimensions behave in the same way

**DOI:** 10.1101/508069

**Authors:** Amit Yashar, Xiuyun Wu, Jiageng Chen, Marisa Carrasco

## Abstract

Humans often fail to identify a target because of nearby flankers. The nature and stage(s) at which this “crowding” occurs are unclear, and whether crowding operates via a common mechanism across visual dimensions is unknown. Using a dual estimation report, we quantitatively assessed the processing of each feature alone and in conjunction with another feature both within and between dimensions. Crowding emerged due to confusion between orientations or colors of target and flankers, but averaging of their spatial frequencies (SFs). Furthermore, crowding of orientation and color were independent, but crowding of orientation and SF were interdependent. This qualitative difference of crowding errors across dimensions revealed a tight link between crowding and ‘illusory conjunctions’ (mis-binding of feature dimensions). These results and a computational model suggest that crowding and illusory conjunction in the visual periphery are due to pooling across a joint coding of orientation and spatial frequencies but not of color.

## Introduction

Recognition of peripheral objects is fundamentally limited by their spacing, not just by the visibility of their features. A target that can be easily identified when presented alone becomes unrecognizable when presented along nearby flankers, i.e. ‘crowded’[1-3]. This breakdown in object-recognition[4] corresponds to the increase in positional uncertainty in the periphery, due to larger receptive-fields[5-7]. Crowding affects various perceptual tasks and hinders the identification of basic stimuli such as letters[2] and faces[8] (see review[9]).

The neural mechanisms underlying crowding are still unknown (reviews by[3,9]). Recent studies have posited that crowding occurs either at an early visual stage, such as V1 or V2 [7,10], or at multiple stages of visual processing and object representation [11-13]. Proposed models explain crowding as reflecting either substitution of objects [14-18] or pooling of features [7,19-23]. Substitution models predict confusion errors, i.e. miss-reporting flankers items instead of the target [14,18]. Pooling models typically predict features averaging errors, i.e. reporting a combination of target and flanker features [23], but recent pooling models attempt to explain both averaging and confusion errors [7,19,21].

Crowding models implicitly assume that the crowding mechanism is in principle the same across various types of stimulus (reviewed by[3,9]); and therefore each proposed model presumably predicts the same type of crowding errors regardless of the types of feature-dimensions, such as orientation, color and spatial frequency (SF). However, whether crowding errors are qualitatively the same across different feature-dimensions and their conjunctions, i.e. feature-binding, is unknown. Most studies have investigated crowding errors within a particular dimension (e.g., orientation [13,14,19,22-25]), and the few investigations of errors within various dimensions could not distinguish averaging and substitution errors, nor could they test for qualitative differences across dimensions [26-28]. Thus, it is still unknown how each of the basic feature-dimensions behaves under crowding and whether they behave interdependently, or independently from each other.

This study had two main goals. First, to test the assumption that the same crowding mechanism applies to different features [9,26]. We investigated the contribution of each flanker to crowding for orientation, SF, and color. Second, we investigated the processing stage at which crowding occurs. Specifically, we asked whether crowding occurs before or after features are bound into an object. To do so, we employed a feature estimation technique that enabled us to simultaneously characterize the pattern of crowding errors within and between feature-dimensions. Thus, we were able to quantitatively assess not only the accuracy of feature perception under crowding conditions, but also the accuracy of feature binding.

## Results

### Experiment 1: Orientation and Color

Observers performed an orientation and color estimation task of a peripheral (7° of eccentricity) colored T-shaped target (**Figure 1A**; see Methods). The target could be either alone –*target-alone condition*– or flanked by two similar T-shaped items –*flanker conditions*. Target and flankers were radially aligned on the horizontal meridian axis, either on the left or right hemifield. The center to center distance between the target and the flankers was 2.1°, within the region of radial crowding [29,30]. The orientation and color of the target were each independently selected at random from two circular parameters spaces. Target orientation was randomly selected out of 180 values evenly distributed between 1°-360°. Target color was randomly selected out of 180 values evenly distributed along a circle in the DKL color space [31,32] Flankers’ orientation and color were selected out of the same circular parameter spaces as the target but differed in values from those of the target. Each flanker had a unique relation to the target in both parameters spaces (negative or positive). This design enabled us to track the direction and distance of estimation errors in both feature-dimensions relative to each flanker. We analyzed the error distributions (the estimated value minus the true target value) of each feature-dimension by fitting probabilistic models (**see Methods**).

**Figure 1.**
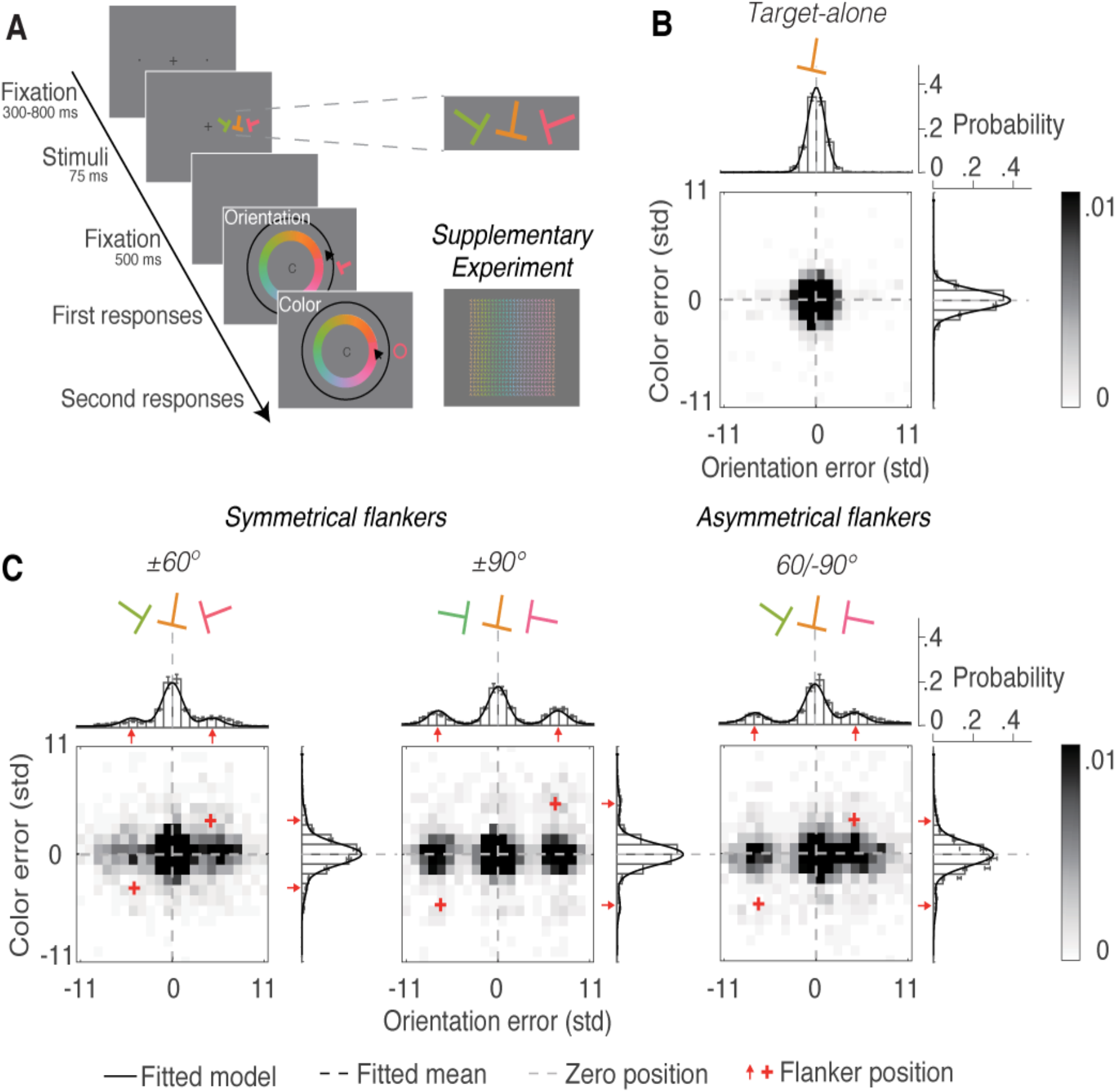
Orientation and color estimation tasks, stimuli, conditions and results in Experiment 1. **A**. Trial sequence. The order of reports (orientation and color) alternated across blocks and was counterbalanced for each observer. The response display In the Supplementary Experiment requires a simultaneous report of color and orientation. Observers had to report both color and orientation by pointing and clicking on the T-shape with the same color and orientation as the target. **B**. Distribution of errors relative to target feature values in the target alone condition. **C**. Distribution of errors in flanker conditions. Error distributions are plotted as a function of the deviation between the estimation report and the target’s feature value for orientation (top), color (right), and their conjunction (heat map). The x-axis is in units of standard deviation (SD) of the error distribution in target-alone trials. Data from asymmetrical flankers were aligned to 60/-90 (as pictured) during analysis. Solid lines indicate the response probabilities predicted by the standard misreport model.

#### Misreporting of orientations or colors

For each feature dimension, the estimation error was defined as the deviation between the reported and the actual feature value of the target (**Figure 1B** and **1C**). We analyzed the error distributions by fitting five probabilistic mixture models to the individual data (**see Methods**). All models included a von Mises (circular) distribution to describe the probability density of precision errors for the target’s feature, and a uniform component to capture the guessing in estimation. We compared between these basic mixture models (models S and B), models that also include a misreports component to describes the probability of reporting one of the flankers to be the target (models SM and BM), and a model that also includes an ‘educated guess’ component to describe the proportion of guessing the target based on flankers values (model EG). We compared models by calculating Akaike Information Criterion with correction (AICc) values for the individual model fits (**Figure 2**). **Table S1** (**Supplementary results**) shows fitted model parameters in each condition and dimension.

**Figure 2.**
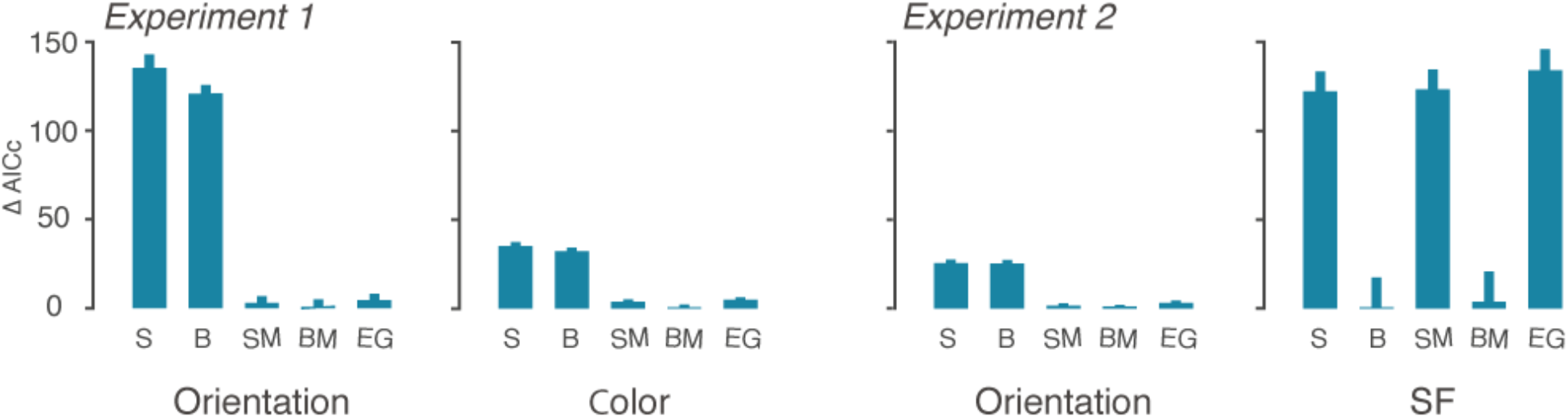
Model comparison. Comparison of individual goodness of fit of the five models to the flanker-conditions data in the orientation and color reports in Experiment 1 and in the orientation and SF reports in Experiment 2. Mean AICc was calculated by subtracting the mean AICc of the best performing model (lowest AICc) from the individual AICc scores of each mixture model. S=standard, B=bias, SM=standard with misreport, BM=bias with misreport, EG = educated guesses. Error bars are 1 SEM.

To compare crowding for orientation and color, we converted each feature dimension values to units of standard deviations (SD) of the error distribution in target-alone trials. So that in each feature dimension target-flanker distance is presented in relation to observer’s precision in the target-alone trials. These standardized units of target-flanker distance (±4.60 and ±6.90 SD for orientation and ±3.43 and ±5.15 SD for color) confirmed that the effect of flanker interference is comparable across feature-dimensions (**Figure 1B** and **1C**).

On target-alone trials, the error distributions for both feature dimensions were well-described by a von Mises distribution centered on the target value with an added non-zero uniform distribution (*γ*) for both orientation and color (**Figure 1B**), *t*(13)>2.98, *p*<.005, indicating that a small yet significant proportion of the responses were statistically unrelated to the target (i.e. guessing).

For both orientation and color, models with misreports components outperformed the models without the misreport component. That is, a significant proportion of errors were centered on the value of each of the two flankers, impairing the fit of a single von Mises distribution to the data. Importantly, adding an ‘educated guess’ component to the misreporting model did not impove the fitting, indicating that observers were unaffected by the partial correlation between target and flankers values. Guessing rate was higher in the flanker conditions, *t*s(13)>2.19, *p*s<.05 (**Table S1**). The misreporting rate (*β*) was larger than zero in all flanker conditions, *t*s(13)>5.22, *p*s<.001. The variability (*α*) of errors centered on the target increased significantly relative to the target-alone condition, *t*s(13)>3.32, *p*s<.01, indicating that crowding led to a reduced precision, and increased the guessing rate and misreporting errors.

Next we compared misreporting rates between orientation and color. Orientations were misreported more than color: orientation averaged *β* =0.27±0.03 vs. color averaged *β*=0.086±0.01, *t*(13)=7.64, *p*<0.001, indicating a larger crowding effect for estimation of orientation than for estimation of color.

#### Observers misreport orientations independently from color

To assess whether orientation and color errors occur before or after orientation and color are bound, we used trial-by-trial correlation between orientation errors and color errors to test the interdependency of crowding errors across feature-dimensions. The joint distributions in each condition are presented in **Figures 1B and 1C** (bottom panels). In the target-alone condition, only 4 out of 14 observers showed a significant Pearson correlation between orientation and color errors, overall mean r=0.06, SE=0.04. In the flanker conditions (all three conditions collapsed), only 3 observers showed a significant correlation, overall mean *r*=0.03, SE=0.02. These findings show that orientation and color estimation were predominantly uncorrelated.

In both feature-dimensions the nature of errors was largely the same: observers reported the orientation or color of a flanker instead of that of the target (misreporting errors). Note that it is unlikely that orientation errors were due to combining target and flanker T-shape parts (e.g. combining target ‘steam’ wit flanker ‘hat’), as such combinations would lead to reporting a vast range of orientations which would be reflected by an increase in the uniform distribution rather than by misreporting errors. The misreporting errors in color could not be explained by optical blur, as such blur predicts averaging errors. Importantly, orientation and color misreporting were independent from each other, suggesting that orientation and color are unbound under crowding conditions.

An alternative explanation is that the separate reports for color and orientation encouraged observers to separately encode color and orientation. Hence, the uncorrelated errors across dimensions could have been due to response strategy rather than unbounded perceptual representation of features. To rule out this alternative explanation, we conducted a control experiment in which observers reported both orientation and color of the target with a single response (**Figure 1A**). This experiment yielded converging results (**Supplementary Experiment and Figure S1 in Supplementary information**): uncorrelated trial-by-trials errors were maintained even when observers simultaneously reported both dimensions. Taken together, these results show that orientation and color crowding errors occur before features are bound into an object.

### Experiment 2: Orientation and Spatial Frequency

In this experiment we tested whether the pattern of results obtained in Experiment 1 would also emerge with different stimuli and feature dimensions. Observers viewed sinusoidal gratings (Gabors) and estimated the target’s spatial frequency and orientation (**Figure 3A**). These two dimensions are jointly coded by individual neurons in V1 [33-35], thus we hypothesized instead of independent, crowding errors may be interdependent.

**Figure 3.**
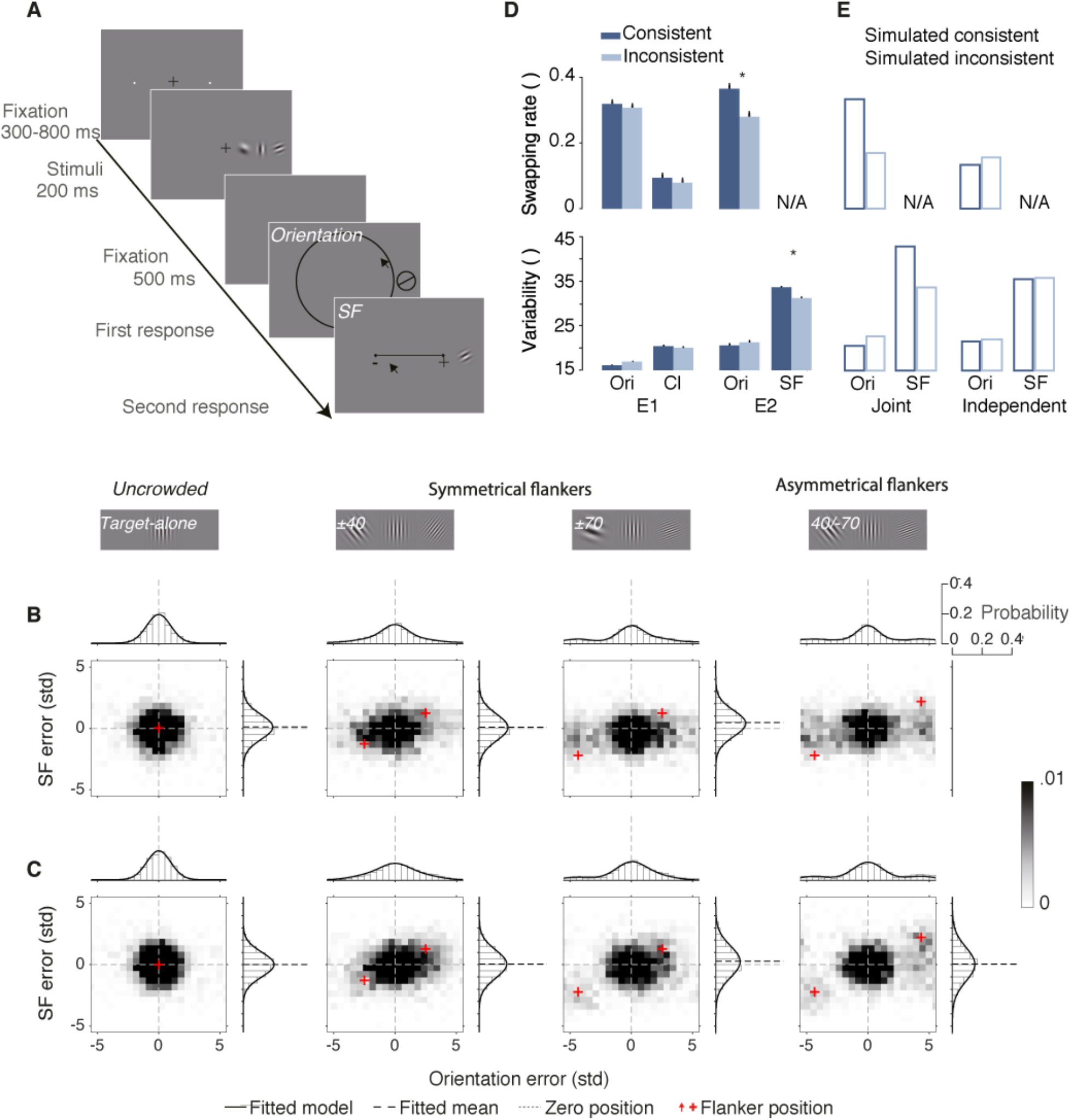
Stimuli and results of Experiment 2, simulated data, and analysis of feature-dimension interdependency in Experiment 1, 2 and simulated data. **A**. Orientation and SF estimation tasks and stimuli in Experiment 2. Illustration of a trial sequence. The order of feature report – SF and orientation – alternated across blocks and was counterbalanced for each observer. **B**. and **C**. Distributions of errors relative to target feature values in Experiment 2 **(B)** and the simulated data for the joint coding model **(C)**. For each condition, error distributions are plotted as a function of the deviation between estimation report and target feature values for orientation (top), SF (right), and their conjunction (heat map. Solid lines are model fits for orientation (standard misreport model) and SF (bias model). The model simulation produced the same distinctive pattern of crowding errors for orientation and SF: misreporting errors for orientation, and increases in variability and mean bias in 40/-70 flanker conditions for SF. **D**. and **E**. Comparisons of misreport rates (top panels) and SD (bottom panels) between consistent and inconsistent trials in Experiments 1 and Experiment 2 **(D)**, and in the simulated data **(E)** of the Joint coding model and independent distribution model. The Joint coding simulated model showed the same pattern of consistent effect as Experiment 2. Error bars are 1 SEM [36]. Asterisks indicate p<.01

#### Misreporting of orientations but averaging of spatial frequencies

Error distributions differed between orientation and SF. Whereas orientation errors were best-described by misreporting models, SF errors were best-described with a single Gaussian function with an added uniform distribution (bias models) (**Figure 2**). In both feature-dimensions the best-fitted models outperformed the ‘educated guess’ model. **Table S1** shows best-fitting model parameters for each condition and dimension.

##### Orientation

As in Experiment 1, significant proportions of orientation error in the flanker conditions were due to misreporting, *t*s(13)>3.6, *p*s<.004, and guessing, *t*s(13)>3.6, *p*s<.003 (Figure 3B). But, the variability of the von Mises distribution between flanker and target-alone conditions were equivalent, *t*s<2.1, *p*s>.06. These results indicate that orientation estimation errors were due to increases in guessing and misreporting rate.

##### SF

The variance of SF errors was larger in the flanker conditions than the target-alone condition (Table S1), *t*s(13) > 3.9, *p*s < .002. The proportion of guesses did not increase compared to the target-alone condition, *t*s(13)<2.1, *p*s>.05. The mean (*μ*) of the Gaussian distribution in the 40/-70 condition was significantly biased, *t*(13)=4.69, *p*< .001, toward the -70 flanker (negative bias) compared to the target-alone condition within the same target SF range (**Table S1**). No effects of bias were found in ±40 and ±70 conditions compared to the target-alone condition, *t*s(13)<0.8, *p*s>.4.

#### Observers misreport orientations interdependently of averaging of spatial frequencies

Heat maps of the joint distributions in each condition are presented in Figure 3B. Orientation and SF estimation errors were uncorrelated in the target-alone condition, only one observer showed a significant linear correlation; mean *r*=0.002, *SE*=0.02. However, across flanker conditions there was a significant linear correlation between orientation and SF errors, *r*=0.138, *SE*=0.03, *p*s < .0001, indicating that orientation and SF crowding errors were interdependent. Individual trial-by-trial linear correlations showed significant (*p*s < .04) correlation in 10 out of 14 observers (*Mean r*=0.14, *SE*=0.03).

To further assess the interdependency of crowding errors across features-dimensions and to compare the interdependency in Experiments 1 and 2, we analyzed the distribution of errors in one dimension based on observers’ errors in the other dimension. That is, we tested whether the direction (with respect to flanker values) of an error in one feature-dimension was dependent on the direction of the error in the other feature-dimension (see Method). To this aim, trials were divided into two groups: trials in which estimation errors for both features (orientation vs. color or SF) were toward the same flanker (*consistent trials*), and trials in which errors for each feature were towards separate flankers (*inconsistent trials*). **Figure 3D** plots the effects of consistent vs. inconsistent errors across dimension on misreporting rates and estimation variability for color (Experiment 1), orientation (Experiment 1 and 2), and SF (Experiment 2). The effect of consistency was only found between orientation and SF but not between orientation and color (see **Supplementary Results** for detailed results).

#### Simulation of pooling of a joint population coding can explain orientation and spatial frequency crowding

To test whether the results of Experiment 2 can be explained by a representation in which orientations and SFs are bound, we fitted a biologically plausible computational model that simulates a population of neurons. In the *joint coding* model the populations jointly code orientations and SFs (**Figure 4**, and **Supplementary Method and Table S2**), whereas in the *independent coding* model the populations separately code orientations and SFs (**Table S3**). The models relies on population coding principles and explains crowding as a weighted sum of target and flanker feature values within a receptive field [19,22]. We used a single bivariate Gaussian distribution to simulate the joint coding of orientation and SF and two univariate normal distributions to simulate the separate representation of orientation and color. To explain averaging in SF reports and in orientation reports, the model assumes pooling over a larger region of space for SF than orientation. This assumption is related to the fact that SF judgments involve assessing the width and distance of multiple cycles inside the Gabor [37-40], whereas the orientation judgment may involve assessing a more narrow region of space, e.g. a region with higher aspect ratio [41]. In the simulation, coding precisions (the inverse of the SD of the Gaussian response function) and the SD of the Gaussian receptive field, were directly responsible for orientation estimation variability and misreporting rate, respectively. Therefore, these parameters were determined based on the fitting of he probabilistic standard misreport model to the observed orientation data. Both simulated models were fitted to the results of Experiment 2 (both *r*^2^ = 0.88). However, unlike Experiment 2 the independent coding model showed no correlation between orientation and SF (mean *r* = 0), and no effect of consistency (**Figure 3E**). The joint coding model, on the other hand, showed the same interdependent pattern as the results of Experiment 2 (**Figure 3C**), including the correlation between orientation and SF (mean *r* = 0.23), and the increase in misreporting rate of orientations and SD of SF in consistent than inconsistent trials (**Figure 3E**). These results support the conclusion that joint coding of orientation and SF underlies the results of Experiment 2.

**Figure 4.**
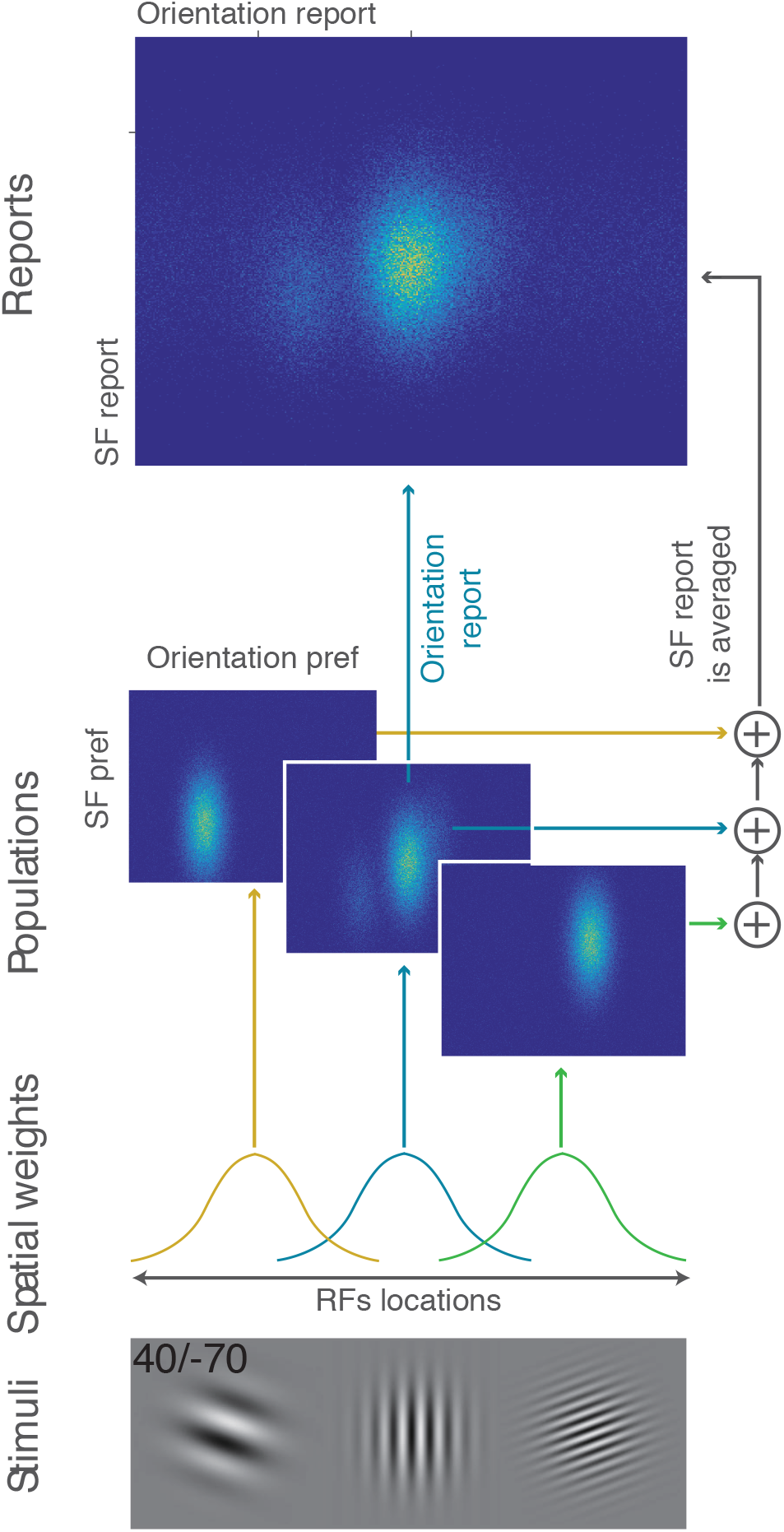
Simulation of population neural activity that jointly codes for orientation and SF fitted to Experiment 2. The model simulates the firing rates of three populations of neurons with receptive field (RF) locations, orientation preference, and SF preference. Stimuli (Bottom row): A target presented with two flankers (asymmetrical flanker condition 40/-70). Spatial weights (second row from bottom): Normalized Gaussian function determines the population level to a stimulus as a function of its relative distance (compared to other stimuli) from the center of RF. Population (third row from bottom): A bivariate probability function with orientation preference (horizontal) and SF preference (vertical). Orientation and SF arrangement is centered over the target orientation and SF values. Reports (top panel): Report of orientation is based on a single population over the target location. Report of SF is based on pooling over different locations, i.e. pooling over three population with RF centered over each location).

## Discussion

In this study we simultaneously characterized the pattern of crowding errors within and between feature-dimensions. In three experiments we demonstrate variations in the pattern of crowding errors based on the specific feature dimension (orientation, color & SF) and their conjunctions. Crowding is more pronounced for orientation than for color. Crowding reflects misreporting a flanker orientation or color instead of those of the target, but averaging of their SFs. The pattern of results was contingent on the feature dimension type but not the stimulus type: observers misreported the target orientation regardless of whether the stimulus was a colored T or a grating. The distinct pattern of crowding errors in each dimension suggests a distinct representation for each feature dimension.

Crowding errors for orientation and SF were interdependent, but those for orientation and color were independent. These findings were shown by the analysis of the joint distribution of feature-dimension errors and trial-by-trial correlations. Moreover, comparison of model parameters revealed higher orientation misreporting rates when SF and orientation errors were toward the same flanker, but that was not the case for orientation and color. These findings suggest that the spatial integration that underlies crowding operates after orientation is bound with SF but before it is bound with color.

### Not all features behave the same under crowding: errors within feature-dimensions

The present study challenges many models’ implicit assumption that crowding operates in the same manner across different feature-dimensions (reviewed by[3,9]). Investigations supporting this view had not tested for qualitative differences in the pattern of errors across dimensions [26-28]. Here, within the same display we show both quantitative and qualitative differences in the pattern of errors across different dimensions. First, orientation misreporting is three times more likely than color misreporting, demonstrating that estimation of orientation is more susceptible to crowding than estimation of color. Second, whereas orientation or color was misreported, SFs were averaged in the SF dimension. This variation between misreporting and averaging occurred within the same stimuli and display (Gabor). It has been proposed that observed errors may be contingent on the target-flankers orientation distance [19,42] (but see[14]). Here, variation between misreporting errors and averaging errors cannot be explained by target-flanker distance. Observers misreported orientation and averaged SF even when the distance in feature-space was effectively the same (**Figure S2**). Thus, this study shows that crowding varies as a function of the feature-dimension being reported.

### Crowding and feature-binding: errors between feature-dimensions

Numerous studies demonstrating observers’ failure to correctly report the conjunction of feature-dimensions of peripheral items (e.g. color and orientation), attributed this phenomenon known as ‘illusory conjunction’ to limited attention (for review see [43]). Alternatively, it has been proposed that ‘illusory conjunctions’ could be due to crowding [1,4,21,27] or to averaging of features [44]. However, studies that investigated crowding and binding have yielded inconsistent results [27,28]. Crowding has been reported to occur either before [27] or after [28] features are bound.

In this study we quantitatively assessed, for the first time, crowding errors both within and between feature-dimensions. The results reveal that under a crowded display color and orientation remain unbound, even when both dimensions are jointly reported. Unlike color, SF remains bound with orientation under crowding displays; observers tended to misreport flanker orientation and average SF with the same flanker. This contingency of binding errors on the specific feature-dimensions may explain why some studies using a set of stimuli suggested that crowding reflects interference in feature-binding [4,27], whereas another study using a different set of stimuli, suggested that crowding follow feature-binding [28].

## Conclusions

This study directly links two mostly-independent topics of research –crowding and feature binding– and challenges conventional views in each of them. By testing crowding both within and between feature-dimensions we show that it is not a uniform phenomenon: it reflects different operations depending on the specific feature-dimensions and their conjunctions. Both the data analysis and our model simulation suggest that crowding reflects spatial integration over neural populations that encode both orientation and SF together but color separately.

## Acknowledgments

This study was supported by The Israel Science Foundation (grant No.111/15) to AY, and NIH-Grant R01 EY016200 to MC. We would like to thank Denis Pelli and members of the Carrasco lab for their useful comments.

## Methods

### Experiment 1: Orientation and color

#### Observers

Fourteen undergraduate and graduate students from New York University participated in this experiment (five females, age range 18 to 29 years old, 21.00 ± 3.28). All observers were naïve to the purposes of the study, and all reported having normal or corrected-to-normal visual acuity and normal color vision. Written informed consent was obtained from all participants before the experiment. The University Committee on Activities Involving Human Subjects at New York University approved experimental procedures.

#### Apparatus

Stimuli were programmed in Matlab (The MathWorks, Inc., Natick, MA) with the Psychophysics Toolbox extensions [45] and presented on a gamma-corrected 21-in CRT monitor (Sony GDM-5402, with 1280 × 960 resolution and 85-Hz refresh rate) connected with an IMac. A chin-rest set the 57-cm viewing distance. Colors and luminance were calibrated using SpectraScan Spectroradiometer PR-670 (Photo Research) spectrometer. Eye movements were monitored and recorded by an Eyelink 1000 (SR Research) infrared eye tracker. Observers used the mouse to generate responses.

#### Stimuli and procedure

**Figure 1A** illustrates a trial sequence. Each trial began with the presentation of a centered black plus (+) sign (fixation mark) subtending 0.5°, along with two dots subtending 0.05° in radius. The dots were presented on the horizontal meridian one in the left and the other in the right hemifields, each centered at an eccentricity of 7° and indicated the two possible target locations. Following observer fixation (for a random duration between 300-800 ms) the stimulus display appeared for 75 ms. In the stimulus display the target was a T-shaped item, subtending 1.6° X 1.6° and drawn with a 0.3° stroke, presented on the horizontal meridian in either the left or right (randomly selected) hemifields, centered at 7° eccentricity. The orientation and color of the target were each independently selected at random from two circular parameters spaces. Target orientation was randomly selected out of 180 values evenly distributed between 1°-360°. Target color was randomly selected out of 180 values evenly distributed along a circle in the DKL color space [31,32] (See Supplementary method). Stimuli color and background were equiluminant (56 cd/m2).

The target could be either alone – *target-alone condition* – or flanked by two similar T-shaped items – *flanker conditions* – centered on the horizontal meridian to the left and to the right of the target (each 2.1° of center to center distance from the target).

The stimulus display was followed by a blank interval of 500 ms, which was then followed by the response displays that appeared until the observer completed both responses. The response displays included an “orientation circle” (a black circle 0.08° thick with an inner radius of 3.8° around the center of the screen) along with a color wheel (1.5 ° thick and had an inner radius of 2.25°) containing the 180 colors. Observers were asked to estimate the target orientation by pointing and clicking the mouse cursor at a position on the orientation wheel, and to estimate the target color by pointing and clicking the mouse cursor at a position on the color wheel. During report a visual feedback of the selected feature was presented at the location of the target. A letter at fixation (either an O or a C, which stands for orientation first, and color first, respectively) indicated report order, which was counterbalanced across blocks.

##### Phase 1

To test the appropriateness of the estimation task with the equiluminant DKL colors, we tested the target-alone condition. **Figure S3** illustrates the color estimation values as a function of target color values for the selected eccentricity. The results showed a linear relation between estimation and target color values, ruling out the possibility that the task was mediated by color categories (e.g., red, blue).

##### Phase 2

###### Design

There were four conditions: three flanker conditions (**Figure 1C** [Top panels]; ±60, 60/-90 and ±90) and the target-alone condition (**Figure 1B**). Flankers’ orientation and color differed from those of the target by either ±60° or ±90° in orientation space and DKL color space. Each flanker had the same absolute target-flanker difference in both feature-dimensions. Within each feature-dimension, one flanker had a positive and the other negative target-flanker difference (randomly selected), such that each flanker had a unique relation to the target (negative or positive) in both feature-dimensions. This design enabled us to track the direction of estimation errors related to each flanker in both feature-dimensions. The three flanker conditions were based on the combinations of the target-flanker differences with the two flankers: (1) 60 and -60; (2) 90 and -90; (3) 60 and -90, or 90 and -60, which were labeled ±60, ±90 and 60/-90 respectively. Each condition had 200 trials (800 trials overall). Each observer completed 10 blocks of 80 trials over two 50-minute sessions (five blocks per session). In each block there were 20 trials from each of the four conditions. Response display order was counterbalanced across blocks. The experiment began with an 80-trial practice block. Observers were encouraged to take a short rest between blocks.

###### Models and analyses

We analyzed the error distributions by fitting probabilistic mixture models, which were developed from the standard model [46] and the standard with misreporting model [47]: For each trial we calculated the estimation error for orientation and color by subtracting the estimation value from the true value of the target. In the flanker conditions we aligned the data such that across feature-dimensions one flanker was consistently positive, and the other was consistently negative. In the 60/-90 flanker condition we aligned all data to be 60 and -90. These alignments enabled us to track the effect of each individual flanker on the error distribution of each feature-dimension separately and in conjunction with the other feature dimension. We compared among five models:

A. *The standard mixture model* (**Equation 1**) uses a von Mises (circular) distribution to describe the probability density of the pooling estimation of the target’s feature, and a uniform component to reflect the guessing in estimation. The model has two free parameters:

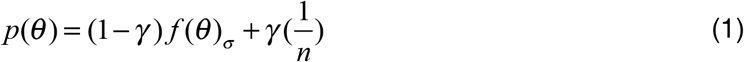

where *θ* is the value of the estimation error, *γ* is the proportion of trials in which participants are randomly guessing (guessing rate), and *f*(*θ*)_*σ*_ is the von Mises distribution with a standard deviation *σ* (variability; the mean is set to zero), and *n* is the total number of possible values for the target’s feature.
B. *The bias mixture model* (**Equation 2**) has three free parameters. In addition to the variability and guessing rate, this model adds a free parameter for the mean (*μ*) of the error distribution:

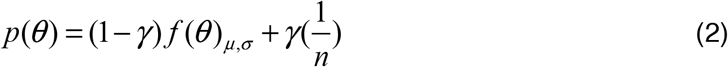
C. *The standard misreport model* (**Equation 3**) has three free parameters. The model adds a misreporting component to the standard mixture model, which describes the probability of reporting one of the flankers to be the target:

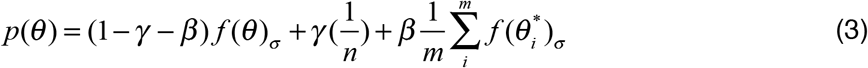

where *β* is the probability of reporting a flanker as the target, *m* is the total number of non-target items (two in the present study), and 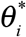 is the error to the feature of the *i*^*th*^ flanker. Notice that *f*(*θ*), the von Mises distribution (for color and orientation) or the Gaussian distribution (for SF) of the estimation errors here describes the distribution when the observer correctly estimated the target’s feature, thus its mean is zero; whereas for 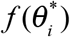 the distribution of estimating one flanker, the mean would be the feature distance of the corresponding flanker to that of the target. The variability of the distributions for each stimulus was assumed to be the same.
D. *The bias misreport model* (**Equation 4**) has four free parameters. In addition to the standard misreport model, there is a free parameter for the mean (*μ*) of the distribution of estimating the target’s feature, to better account for possible pooling and substitution:

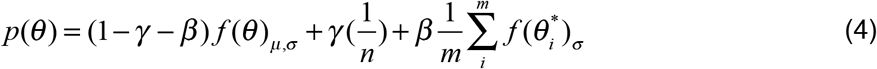
E. *The educated guess model* (**Equation 5**) has four free parameters. The model adds a misreporting component of the guessed stimuli other than the stimuli presented to the standard misreport model. This misreporting component is similar as the misreporting component of the flankers, but has a different probability:

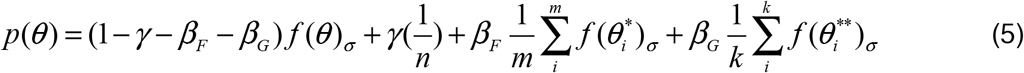

where *β*_*F*_ is the probability of misreporting a flanker as the target, and *β*_*G*_ is the probability of misreporting a guessed feature other than the flankers presented. This model follows the assumption that the participant may have the information of one feature, then guess the feature of the target based on all possible target-flanker distances. This “educated guess” could result in misreporting one flanker as the target, or misreporting features not presented in the corresponding trial. For example, there are four possible target-flanker distances: -90, -60, 60, and 90. In a 60/-60 trial, for each detected feature, there are five possible feature values (including the stimulus itself) that could be the possible target; therefore, there will be two flankers (-60, 60; *m* = 2) with the misreporting probability *β*_*F*_, and ten not presented guessed stimuli (some of them are overlapped, and the non-overlapping feature values are -150, -120, -90, -30, 30, 90, 120, 150; *k* = 10) with the misreporting probability *β*_*G*_.

We used the MemToolBox [48] for fitting the models, and compared Akaike information criterion with correction (AICc) to assess model fits.

### Experiment 2: Orientation and Spatial Frequency

#### Observers

Fourteen undergraduate students from New York University participated in this experiment (12 females, age ranged from 18 to 21 years old, 19.43 ± 0.85). All observers were naïve to the purposes of the study, and all reported having normal or corrected-to-normal visual acuity. Written informed consent was obtained from all participants before the experiment. The University Committee on Activities Involving Human Subjects at New York University approved experimental procedures.

#### Apparatus, stimuli, procedure and design

Sequence of events within a trial and sample stimulus displays are presented in **Figure 3**. The apparatus, stimuli, procedure and design were the same as in Experiment 1 except for the following changes: Target and flankers were sinusoidal gratings (Gabors) with a 2D Gaussian spatial envelope (standard deviation 0.325° and 85% contrast). The orientation and SF of the target were each randomly and independently selected from two parameters spaces. The viewing distance was 91 cm. Target and flankers’ center-to-center distance was 2.15°. The orientation parameter space corresponded to range of angle 1°-180°. Stimulus display duration was 200 ms.

##### Phase 1

To determine the SF values of the estimation task, we tested the target-alone condition with different SF values, linearly spaced. Based on this test we set the SF parameter space to correspond to the range of 1-5 cpd. Because SF discriminability varies across SF values [49], we scaled the 180 unit steps of SF that were used in Phase 2 according to the variation in the estimation task of SF values in Phase 1. To this aim we fit an exponential function to the SD of the estimation data for each SF value in Phase 1 (**Figure S4**).

##### Phase 2

###### Design

Flankers’ orientation and SF differed from the target by either ±40 or ±70 units of orientation and SF units. Each flanker had the same absolute target-flanker difference in both feature-dimensions. Within each feature dimension, one flanker had a positive and the other negative target-flanker difference (randomly selected), such that each flanker had a unique relation to the target (negative or positive) in both feature-dimensions. Target flanker distance within the parameter space was: (1) 40/-40; (3) 70/-70, (3) 40/-70, 70/-40, which were labeled ±40, ±70 and 40/-70 respectively.

In all trials the target orientation was randomly selected between 1°-180° in a step size of 2°. In target-alone trials, the target SF was randomly selected out of the 180 steps, as determined in Phase 1. However, because SF is not a circular space, in the crowding display flankers’ SF values restricted the range of target SF values, such that the target SF ranged from 41-140, 71-110, and 41-110/71-140 SF steps in ±40, ±70 and 40/-70 flankers conditions respectively.

There were two response displays. In the orientation response display observers had to estimate the target orientation by pointing and clicking the mouse cursor at a position on the orientation wheel. In the SF response display observers estimated SF by pointing and clicking the mouse cursor on a centered horizontal line (0.08°-thick two-directional arrow with 11.2° in length). A “-” on one side and “+” on the other side indicated the direction of SF increase. Because SF is not circular, we extended the range of the SF response (0.48-7.35 cpd) beyond that of the target (1-5 cpd).

###### Models and analyses

Because SF is not circular, in the mixture models we used a Gaussian distribution when fitting to SF and von Mises distribution when fitting to orientation. Because in flanker conditions we had to restrict the range of target SF, when comparing SF flanker and the target-alone conditions we equated the range of target SF.

---

**Table S1.**
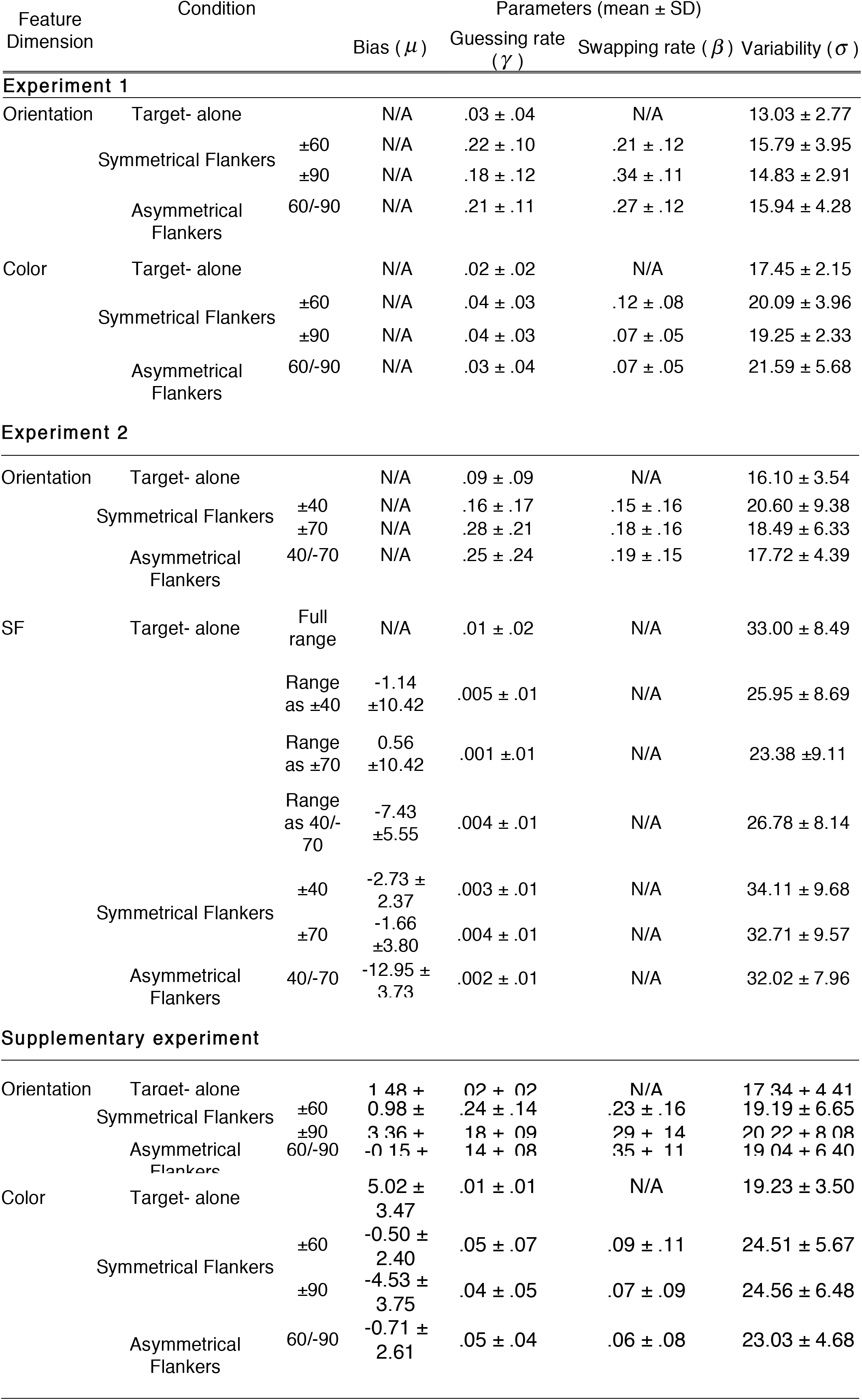
Best-fitting parameters for each condition in Experiment 1, Experiment 2 and Supplementary experiment. For orientation and color, parameters are from the standard mixture model for the target-alone condition, and the standard with misreport model for the flanker conditions. For SF data, parameters are from the bias model.

## Supplementary Experiment

In Experiment 1 and 2 observers reported each feature separately, which may have encouraged observers to decode each feature independently during response, which may have led to estimation errors that are uncorreleated. We addressed this alternative explanation in this Supplementary Experiment by using only one response display and having observers simultaneously report both color and orientation.

## Method

### Observers

Fourteen undergraduate and graduate students from New York University participated in this study (10 females, age range 20 to 27 years old, 21.79 ± 2.04). All observers were naïve as to the purposes of the study, and all reported having normal or corrected-to-normal visual acuity and normal color vision. Written informed consent were obtained from all observers before the experiment. The University Committee on Activities Involving Human Subjects at New York University approved experimental procedures.

### Stimuli and procedure

The stimuli and procedure were the same as in Experiment 1 except for the following changes. Target orientation was randomly selected out of 24 values evenly distributed between 1° - 360°. Target color was randomly selected out of 24 values evenly distributed along the circle in the DKL color space as defined in Experiment 1. The sets of 24 orientations and 24 colors were independently selected randomly for each observer. The response display was a 24X24 matrix of T-shape items, which comprised all the possible conjunctions of target orientation and color (**Figure S1**), and were ordered according to their orientation (vertical order) and color (horizontal order) values. Observers were instructed to report the target by pointing and clicking the mouse cursor over the item with the orientation and color of the target. To facilitate report, the matrix was the same for each observer, and observers were allowed to saccade during the response.

## Results

### Distributions of errors within each feature-dimension

**Figure S1a** plots the distribution of estimation errors for orientation and color in target-alone trials. In both feature-dimensions the error distribution was accurately described by a von Mises function with an added uniform distribution. One participant was excluded from the analysis due to a large variability in orientation errors and poor fits for all models. For orientation errors, the von Mises function centered on 1.48 ± 2.15 away from target value with a variability (*σ*) of 17.34 ± 4.41 could describe well the error distributions, and for color errors the circular Gaussian function centered on 5.02 ± 3.47 away from target value, with a variability of 19.23 ± 3.50, could describe well the error distributions. The probability of the uniform distribution was significantly higher than zero for orientation (*γ*_*c*_ = 0.02 ± 0.02, *t*(12) = 3.43, *p* < .01), and marginally so for color (*γ*_*o*_ = 0.01 ± 0.01, *t*(12) = 1.90, *p* = .08).

**Figure S1.**
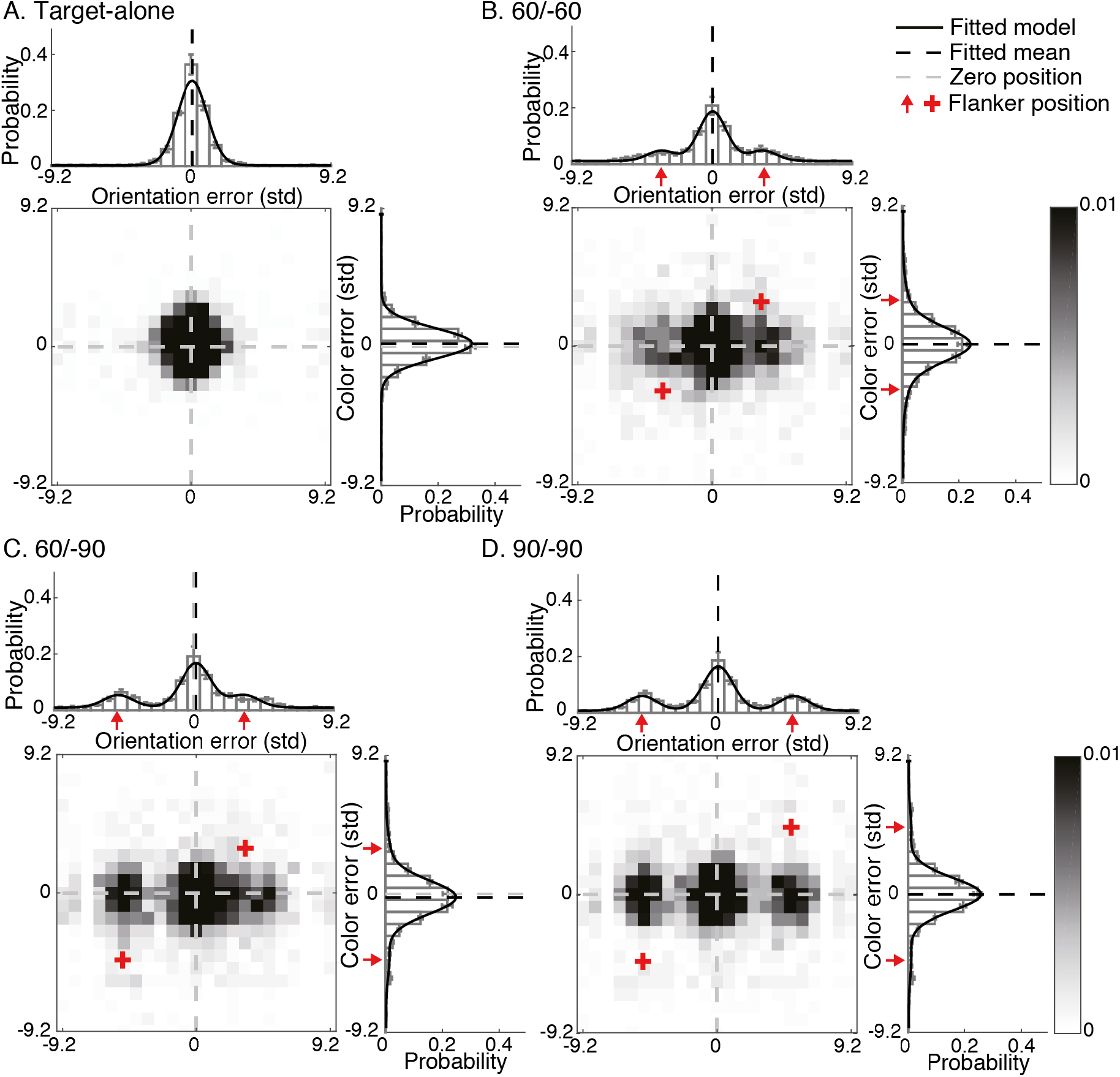
Supplementary Experiment distribution of errors relative to target feature values. Frequency of response is plotted as a function of the deviation between estimation report and target feature values (x-axis units are SD in target-alone trials): for orientation (top), color (right), and conjunction of both features (heat map) in the **(A)** target-alone condition, **(B)** ±60, **(C)** 60/-90 and **(D)** ±90 flankers conditions. Solid lines indicate the response probabilities predicted by the bias with misreport model.

**Figure S1b, S1c, S1d** plot the distribution of estimation errors for orientation and color in ±60, 60/-90 and ±90 flanker conditions respectively. As in Experiment 1 for both orientation and color the models with misreport outperformed the models without the misreport component suggesting that in both feature dimensions observers confused flanker feature with that of the target. We then analyzed the parameters in the individual fitting of the bias with misreport model between conditions. Compared to the target-alone condition, guessing rate was higher in the crowded conditions as indicated by the increase in the uniform distribution, *t*s(12) > 2.73, *p*s< .02. Also, a significant proportion of the errors concentrated (i.e. von Mises distributed) around the value of one of the two flanker items (misreporting errors). The misreporting rate was significantly larger than zero in all flanker conditions, *ts*(12) > 2.60, *p*s<.03. In all flanker conditions, the variability of the color errors concentrated around the target or the flanker showed significant increase compared to the target-alone condition, *t*s(12) > 4.59, *p*s< .001, while the variability of the orientation errors showed no significant increase in ±60 and 60/-90 conditions, *t*s(12) < 2.06, *p*s > .06, but was significantly increased in the ±90 condition, *t* (12) = 2.23, *p* < .05. For color errors, when compared to the target-alone condition, all flanker conditions showed significant difference in mean (more toward the negative direction), *t*s(12) > 5.67, *p*s<.001; when compared to zero, however, only in the 60/-90 condition the mean of the von Mises distribution was significantly different (smaller than zero), *t*(12) = -4.35, *p* < .002. For orientation errors, when compared to the target-alone condition, no flanker conditions showed significant difference in mean, *t*s(12) < 1.34, *p*s > .21; when compared to zero, only in the 60/-90 condition the mean of the von Mises distribution was significantly different (larger than zero), *t*(12) = 2.85, *p* < .02.

### Joint distributions of errors

Similar to Experiment 2, only 3 observers had significant Pearson correlation between orientation and color estimations under the target-alone and the flanker conditions (collapsed), overall mean *r* = 0.04 ±0.03 and *r*=0.02 ±0.02, respectively.

**Figure S2.**
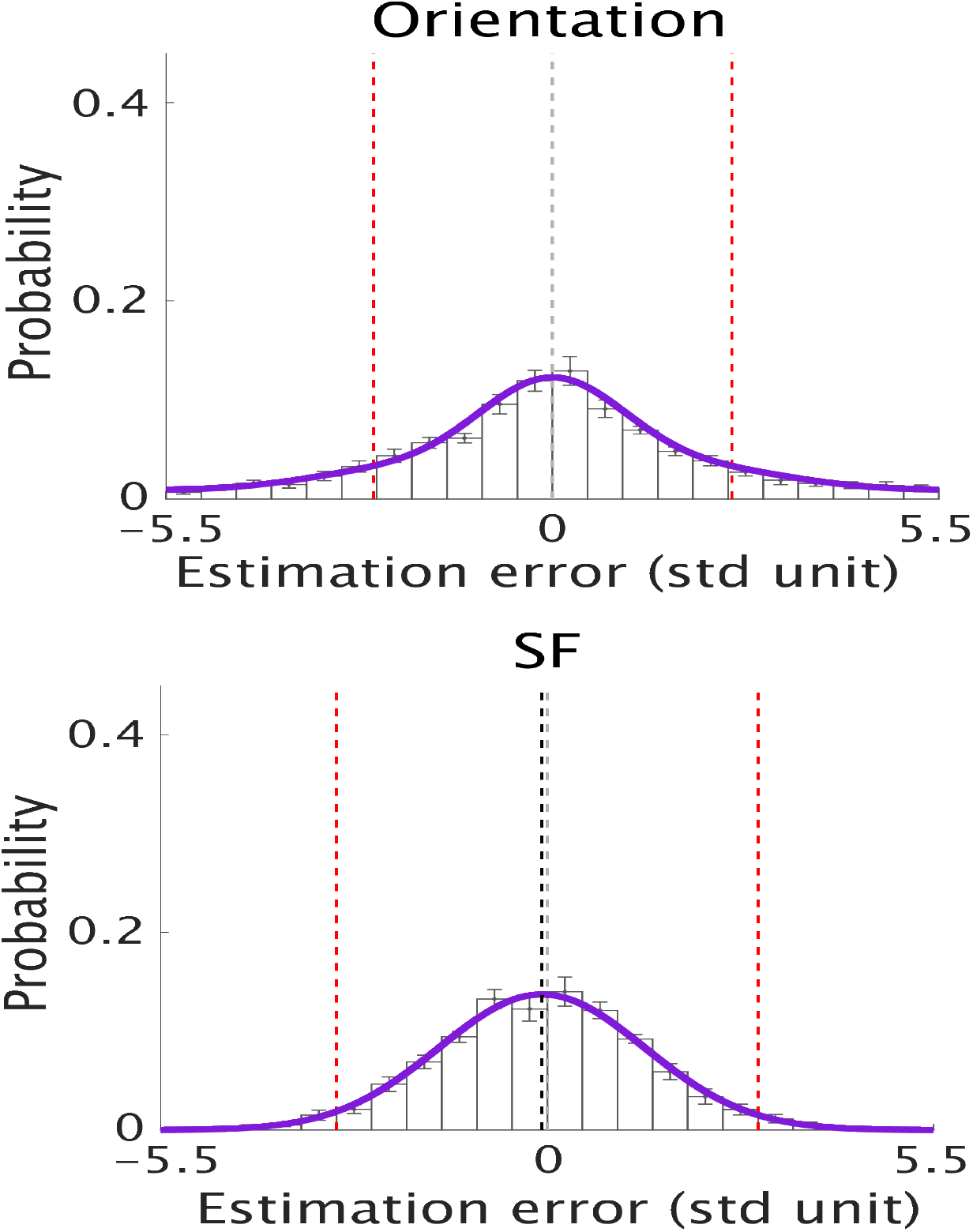
Experiment 2 error distributions for orientation ±40 flanker condition (top panel) and SF ± 70 flanker condition (bottom panel). Even when the flanker feature distance in SD was similar between orientation and SF, observers misreport flanker orientation but averaged SF.

### Supplementary Results

#### Analyses of consistent vs. inconsistent trials

We conducted a two-way (3 flanker conditions x 2 trial types) repeated measures Analysis of Variance (ANOVA) on the parameters of the best fitted models (standard misreport for color and orientation, bias for SF), There were no main effects of consistency on misreporting rate, guessing rate or variability in Experiment 1 in either orientation or color, *Fs*<2.2, *p*s>.1. In Experiment 2, however, the proportion of orientation misreporting errors was higher in consistent than inconsistent trials, *F*(1,13)=11.20, *p*<.01. There was no significant effect of consistency on the variability of errors around the target or guessing rates in orientation, all *p*s >.4. In SF, the variability of errors around the target was higher in consistent than inconsistent trials, *F*(1,13)=19.86, *p*<.002. There was no significant main effect of consistency on mean bias or guessing rate, *p*s>0.07.

## Supplementary Method

### Population coding model

#### Stimulation of population coding of orientation and SF

**Table S2:**
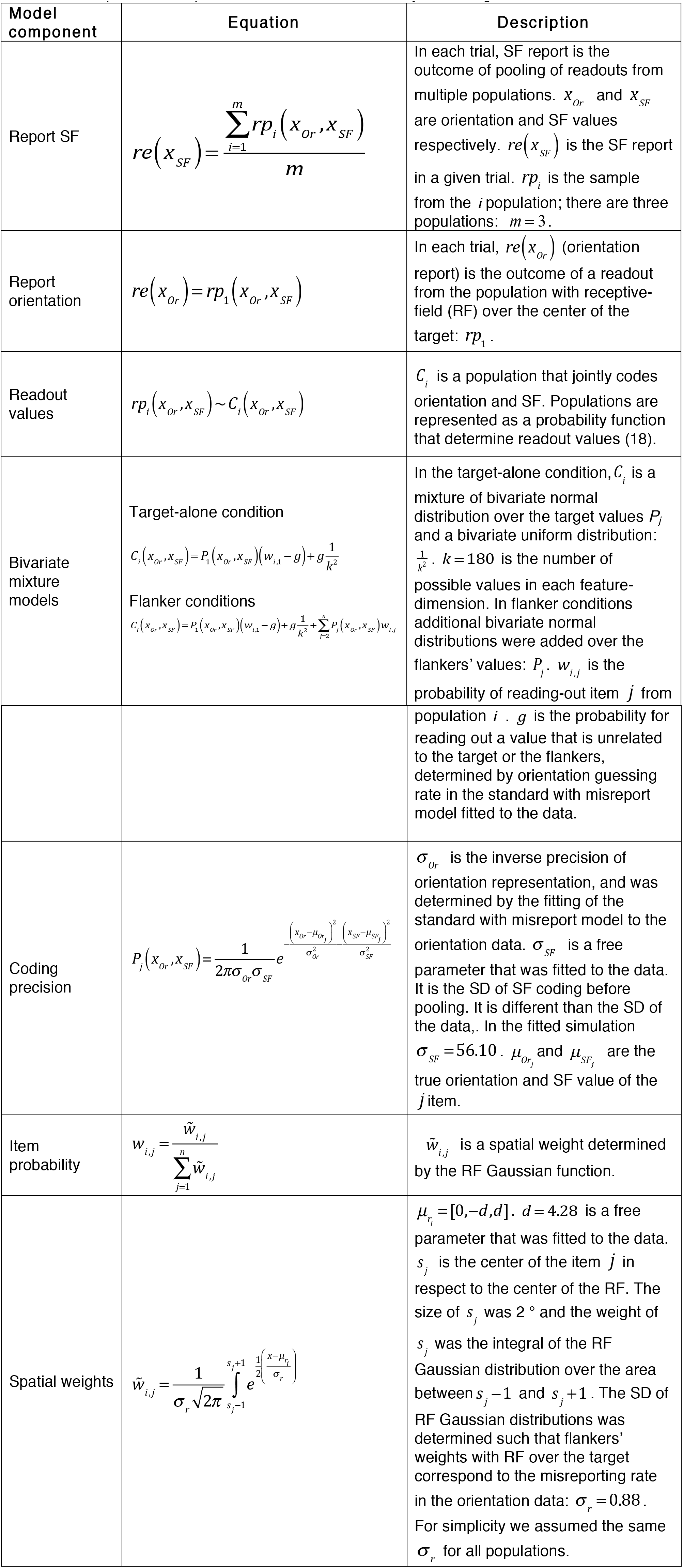
Equations and parameters of the simulation of joint coding of orientation and SF

| Model component | Equation | Description |
| --- | --- | --- |
| Report SF | $re(x_{SF}) = \frac{\sum_{i=1}^m rp_i(x_{Or}, x_{SF})}{m}$ | In each trial, SF report is the outcome of pooling of readouts from multiple populations. $x_{Or}$ and $x_{SF}$ are orientation and SF values respectively. $re(x_{SF})$ is the SF report in a given trial. $rp_i$ is the sample from the $i$ population; there are three populations: $m = 3$. |
| Report orientation | $re(x_{Or}) = rp_1(x_{Or}, x_{SF})$ | In each trial, $re(x_{Or})$ (orientation report) is the outcome of a readout from the population with receptive-field (RF) over the center of the target: $rp_1$. |
| Readout values | $rp_i(x_{Or}, x_{SF}) \sim C_i(x_{Or}, x_{SF})$ | $C_i$ is a population that jointly codes orientation and SF. Populations are represented as a probability function that determine readout values (18). |
| Bivariate mixture models | Target-alone condition<br>$C_i(x_{Or}, x_{SF}) = P_1(x_{Or}, x_{SF}) \left( w_{i,1} - g \right) + g \frac{1}{k^2}$<br>Flanker conditions<br>$C_i(x_{Or}, x_{SF}) = P_1(x_{Or}, x_{SF}) \left( w_{i,1} - g \right) + g \frac{1}{k^2} + \sum_{j=2}^n P_j(x_{Or}, x_{SF}) w_{i,j}$ | In the target-alone condition, $C_i$ is a mixture of bivariate normal distribution over the target values $P_j$ and a bivariate uniform distribution: $\frac{1}{k^2}$. $k = 180$ is the number of possible values in each feature-dimension. In flanker conditions additional bivariate normal distributions were added over the flankers' values: $P_j$. $w_{i,j}$ is the probability of reading-out item $j$ from |
| | | population $i$. $g$ is the probability for reading out a value that is unrelated to the target or the flankers, determined by orientation guessing rate in the standard with misreport model fitted to the data. |
| Coding precision | $P_j(x_{Or}, x_{SF}) = \frac{1}{2\pi\sigma_{Or}\sigma_{SF}} e^{-\frac{(x_{Or}-\mu_{Or_j})^2}{\sigma_{Or}^2} - \frac{(x_{SF}-\mu_{SF_j})^2}{\sigma_{SF}^2}}$ | $\sigma_{Or}$ is the inverse precision of orientation representation, and was determined by the fitting of the standard with misreport model to the orientation data. $\sigma_{SF}$ is a free parameter that was fitted to the data. It is the SD of SF coding before pooling. It is different than the SD of the data,. In the fitted simulation $\sigma_{SF} = 56.10$. $\mu_{Or_j}$ and $\mu_{SF_j}$ are the true orientation and SF value of the $j$ item. |
| Item probability | $w_{i,j} = \frac{\tilde{w}_{i,j}}{\sum_{j=1}^n \tilde{w}_{i,j}}$ | $\tilde{w}_{i,j}$ is a spatial weight determined by the RF Gaussian function. |
| Spatial weights | $\tilde{w}_{i,j} = \frac{1}{\sigma_r \sqrt{2\pi}} \int_{s_j-1}^{s_j+1} e^{\frac{1}{2} \left( \frac{x-\mu_{r_i}}{\sigma_r} \right)^2}$ | $\mu_{r_i} = [0, -d, d]$. $d = 4.28$ is a free parameter that was fitted to the data. $s_j$ is the center of the item $j$ in respect to the center of the RF. The size of $s_j$ was $2^\circ$ and the weight of $s_j$ was the integral of the RF Gaussian distribution over the area between $s_j - 1$ and $s_j + 1$. The SD of RF Gaussian distributions was determined such that flankers' weights with RF over the target correspond to the misreporting rate in the orientation data: $\sigma_r = 0.88$. For simplicity we assumed the same $\sigma_r$ for all populations. |

**Table S3:** Equations and parameters of the simulation of the independent coding of orientation and SF. The model is the same as the joint coding model except for the following changes.

| Model component | Equation | Description |
| --- | --- | --- |
| Readout values | $rp_i(x_{Or}) \sim C_i(x_{Or})$ $rp_i(x_{SF}) \sim C_i(x_{SF})$ | $C_i$ is a population that independently codes orientation or SF. |
| Bivariate mixture models | Target-alone condtion<br>$C_i(x_{Or}) = P_1(x_{Or})(w_{i,1} - g) + g \frac{1}{k^2}$ $C_i(x_{SF}) = P_1(x_{SF})(w_{i,1} - g) + g \frac{1}{k^2}$<br>Flanker conditions<br>$C_i(x_{Or}) = P_1(x_{Or})(w_{i,1} - g) + g \frac{1}{k^2} + \sum_{j=2}^n P_j(x_{Or}) w_{i,j}$ $C_i(x_{SF}) = P_1(x_{SF})(w_{i,1} - g) + g \frac{1}{k^2} + \sum_{j=2}^n P_j(x_{SF}) w_{i,j}$ | In the target-alone condition, $C_i$ is a mixture of normal distributions over the target values $P_j$ and a uniform distribution: $\frac{1}{k}$. In flanker conditions additional normal distributions were added over the flankers' values: |
| Coding precision | $P(x_{Or}) = \frac{1}{\sigma_{Or} \sqrt{2\pi}} e^{-\frac{(x_{Or} - \mu_{Or})^2}{2\sigma_{Or}^2}}$ $P(x_{SF}) = \frac{1}{\sigma_{SF} \sqrt{2\pi}} e^{-\frac{(x_{SF} - \mu_{SF})^2}{2\sigma_{SF}^2}}$ | $\sigma_{Or}$ is the inverse precision of orientation representation, and was determined by the fitting of the standard with misreport model to the orientation data. $\sigma_{SF}$ is a free parameter that was fitted to the data. It is the SD of SF coding before pooling. It is different than the SD of the data,. In the fitted simulation $\sigma_{SF} = 56.10$. $\mu_{Or_j}$ and $\mu_{SF_j}$ are the true orientation and SF value of the $j$ item. |

To fit the model we simulated 2800 trials (same as in the experiment data for all observers together). In each of the four experimental conditions. As in Experiment 2 data, in the target-alone condition we fitted the standard mixture model, and in the flanker conditions we fitted the standard with misreport model to the orientation report and the with bias model to the SF report. To fit the simulations to the data, we minimized the sum of squared errors between the simulated data and Experiment 2 data using the Matlab function ‘patternsearch’. Several starting points for the parameters were used to check the consistency of the model fits. We compared between two

#### DKL colors

The DKL spherical color space is defined by three orthogonal axes: two opponent chromatic cone dimensions (L-M and S-(L+M)) and one luminance (L + M). Here we specified DKL color by calibrating the monitor using the Spectroradiometer PR-670 and the PsychoPy toolbox (http://www.psychopy.org/general/colours.html#dkl1984).

Specifying coordinates in DKL requires selection of a reference cone activation vector, usually derived from the background of the visual display and representing the adaptation state. To simplify the calculations for this experiment, we chose cone values corresponding to the achromatic (white) points at the background luminance (56 cd/m2). Following the convention of Derrington et al. (1984), DKL values were expressed in spherical coordinates. Luminance corresponded to elevation angle, which was zero for all colors here. Azimuth angles controlled hue, with the zero azimuth representing activation along the L - M axis in the direction of positive L-M values. Radius controlled distance from the white point, or saturation. To generate the color wheel we picked 180 hue values (angles), which were evenly distributed between 0°-359°, with a fixed radius of 0.9.

**Figure S3.**
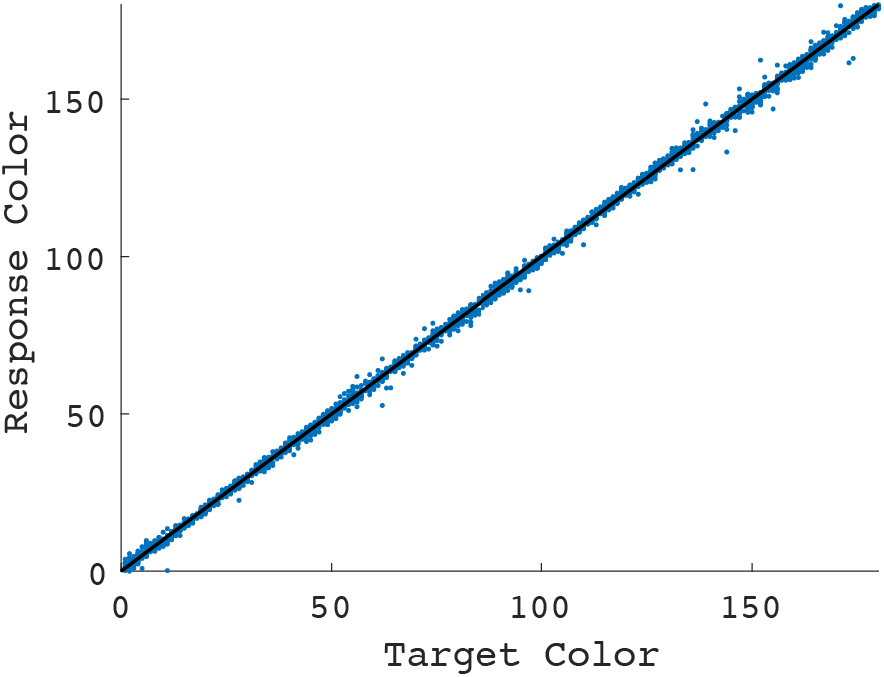
Color estimation performance. Scatter plot of the color estimation (y-axis) as a function of target color values (x-axis) in the target-alone condition. The results show color estimates and the true color value are along the diagonal, indicating that colors were accurately estimated and rules out the possibility that the task could have been mediated by color categories (e.g., red, blue).

**Figure S4.**
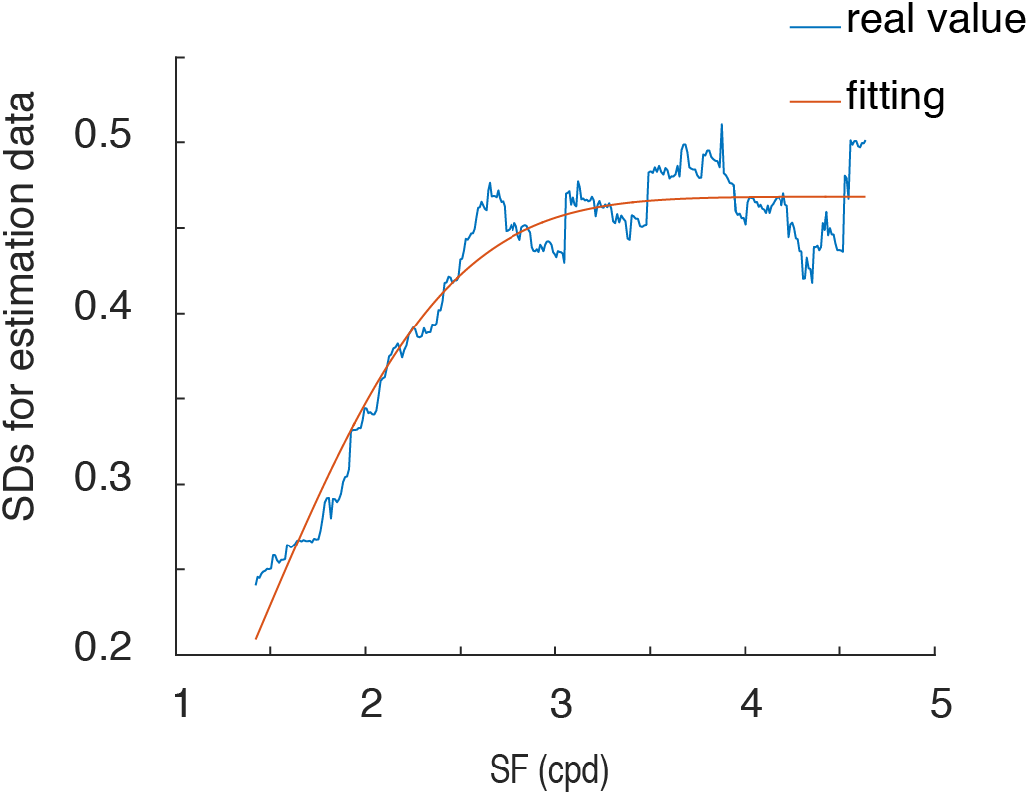
Phase 1 of Experiment 2. SD for the SF estimation task (y-axis) as a function of SF value (x-axis). An exponential function fit the data and was used to scale the SF steps in Phase 2.

